# Mechanical Deformation Behaviors and Structural Properties of Ligated DNA Crystals

**DOI:** 10.1101/2022.06.13.495931

**Authors:** Ruixin Li, Mengxi Zheng, Anirudh Sampath Madhvacharyula, Yancheng Du, Chengde Mao, Jong Hyun Choi

**Affiliations:** School of Mechanical Engineering, Purdue University, West Lafayette, Indiana 47907; Department of Chemistry, Purdue University, West Lafayette, Indiana 47907

**Keywords:** DNA self-assembly, DNA crystal, mechanical property, molecular dynamics simulation, nanoindentation

## Abstract

DNA self-assembly has emerged as a powerful strategy for constructing complex nanostructures. While the mechanics of individual DNA strands have been studied extensively, the deformation behaviors and structural properties of self-assembled architectures are not well understood. This is partly due to the small dimensions and limited experimental methods available. DNA crystals are macroscopic crystalline structures assembled from nanoscale motifs via sticky-end association. The large DNA constructs may thus be an ideal platform to study structural mechanics. Here we have investigated the fundamental mechanical properties and behaviors of ligated DNA crystals made of tensegrity triangular motifs. We performed coarse-grained molecular dynamics simulations and confirmed the results with nanoindentation experiments using atomic force microscopy. We observed various deformation modes including un-tension, linear elasticity, duplex dissociation, and single-stranded component stretch. We found that the mechanical properties of a DNA architecture are correlated with those of its components, however the structure shows complex behaviors which may not be predicted by components alone.

## INTRODUCTION

DNA nanotechnology offers as a powerful bottom-up manufacturing approach with excellent programmability and structural predictability^1,2^. DNA self-assembly can construct intricate structures with sub-nanometer precision using DNA tiles^3-5^, origami^6-8^, or bricks^9-11^. The tile-based method involves several single-stranded DNA (ssDNA) oligonucleotides to form a motif, which then self-assembles into 2D or 3D configurations via sticky-end association. In DNA origami, a long genomic scaffold strand is folded as designed and stabilized with the help of staple oligonucleotides. The DNA brick approach uses many unique ssDNA strands in a single step annealing.

Although various arbitrary nanostructures can be achieved, the related mechanics are still not well understood. This fundamental knowledge is curial for designing complex architectures and transforming them into functional materials. Previous studies reported deformation behaviors and mechanical properties of double-stranded (ds) DNA^12,13^, rods^14,15^, and polyhedrons^16,17^. These studies revealed some mechanical properties (*e.g*., Young’s modulus). However, they are limited to simple structures due to the difficulties in applying external forces on the small dimensions. In complex architectures, it is extremely challenging to apply extension, compression, and twist forces using optical and magnetic tweezers. There are also reports on programmed reconfigurations of DNA structures; however, the loadings were introduced indirectly, most commonly through strand displacement-rehybridization^18-21^ and DNA intercalation^22-25^. Direct mechanical studies to unveil the basic mechanics of complex geometries are lacking.

DNA crystals are macroscopic crystalline structures assembled from nanoscale DNA motifs.^26^ The motifs include tensegrity triangles with three pairs of protruding sticky ends in three directions (total six sticky ends), squares with eight sticky ends,^27^ and hexagons with twelve sticky ends.^28^ The units can self-assemble into a 3D crystal larger than 100 µm via sticky-end association, and their physical properties depend on several parameters including motif design and edge length.^29-31^ Thus, the macroscopic DNA constructs could serve as a platform to study deformation behaviors and structural properties. However, the assemblies are unstable and fragile. They need high ion concentrations (*e.g*., 50 mM Mg^2+^) in the buffer to maintain their shape.^31^ They may be reinforced by triplex formation^32^ or enzymatic ligation^31^. The ligated crystals can survive no ion environment such as deionized (DI) water and cycles between dehydration and rehydration.^31^ Therefore, ligated DNA crystals may be ideal to investigate structural mechanics of complex DNA assemblies. For example, key elastic properties such as Young’s modulus and yield strength as well as plastic behaviors and failure mechanics may be studied. It will also be possible to elucidate the relationship between structural properties of DNA assemblies and nano-motif designs.

Here we have investigated the deformation behaviors and mechanical properties of ligated DNA crystals. Molecular dynamics (MD) simulations were performed based on a coarse-grained model. Given the computational capacity and time, we built a model of a fully ligated DNA crystal made of 5×5×5 triangular motifs with a length of 2 or 4 full turns in each direction (termed 2T and 4T crystals). Tensile forces were applied by immobilizing one surface (bottom) and pulling the other (top). As the top and bottom surfaces moved away from each other (with increased tensile forces), the crystal deformed. We observed various deformation modes with the 4T crystal: un-tension, linear elasticity, dsDNA dissociation, and ssDNA stretch. We extracted force-extension and stress-strain plots from which we analyzed structural properties such as Young’s modulus and yield strength. We found that the plastic deformation is due to the dissociation of dsDNA into ssDNA and the severe contraction of cross-sections. In parallel, we constructed macroscopic crystals from 4T motifs and strengthened them using DNA ligase enzymes (similar to the computer simulations). Nanoindentation experiments were performed using atomic force microscopy (AFM) and probed the elastic range. From the simulation and experiment, we concluded that the 4T crystal has a structural Young’s modulus of about 1 MPa which is two orders of magnitude smaller than that of a component dsDNA. The mechanical behaviors can be modulated by changing the crossover density in the motif. The tightly formed 2T crystal does not show the un-tension regime nor undergo significant cross-sectional changes. The overall properties of a DNA architecture are correlated with those of the component as anticipated, yet the structure clearly demonstrates complex behaviors that may not be predicted by components alone.

## COMPUTATIONAL AND EXPERIMENTAL METHODS

### Design of the DNA Crystal

Figure 1(a) illustrates the tensegrity triangle motif which has three strands in three independent directions, marked in red, green, and blue. A strand in the center (cyan) holds the strands together. To secure the directions, another three strands are placed to combine two neighboring directions together, simultaneously offering sticky ends. The tensegrity triangle motif is symmetric in terms of length and strand arrangement but the sequences can be different (see SI).^30^ In this study, we explored two different edge lengths: 2 and 4 full turns, termed 2T and 4T motifs, respectively. The motifs can self-assemble via sticky-end association (Figure 1(b)), forming 2T or 4T crystals depending on the length.

**Figure 1.**
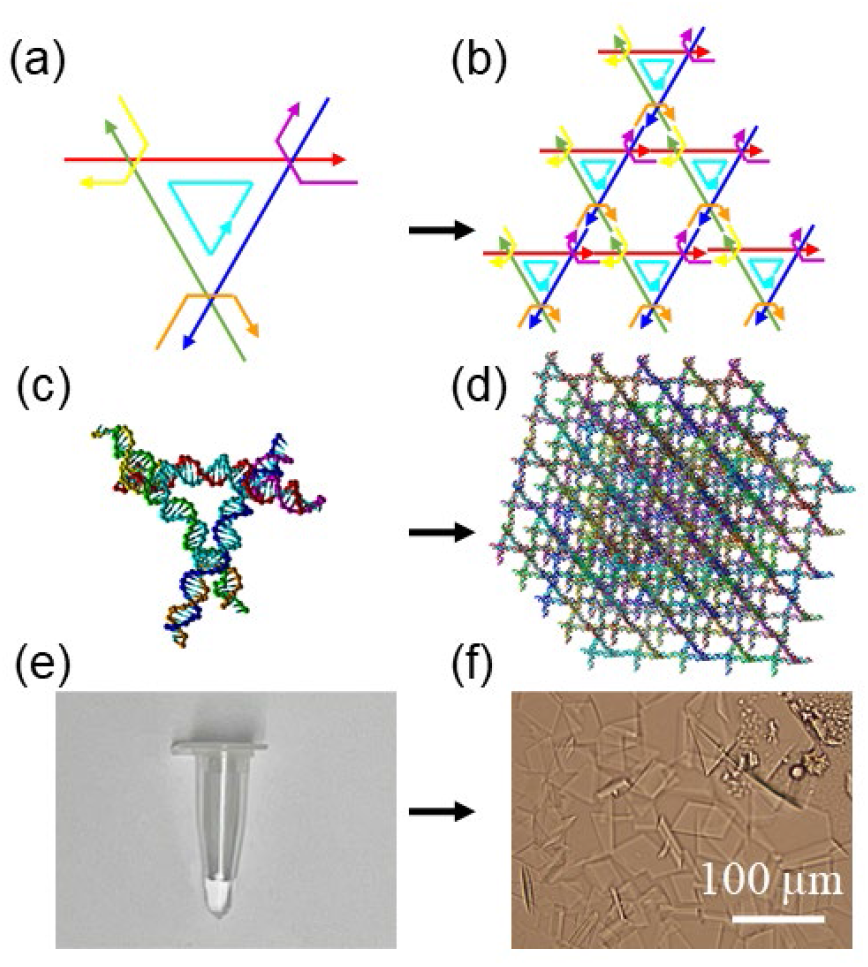
(a) Schematic of a tensegrity triangle motif. Three independent directions are marked as red, green, and blue. The center cyan strand holds the three strands together. Yellow, orange, and purple strands secure the directions with sticky ends. (b) Schematic of the triangle motif assembly via sticky-end association. (c) A tensegrity triangle motif with a length of 4 full turns (4T motif). The molecular model is constructed in the oxDNA software. (d) A DNA crystal made of 5×5×5 4T motifs (4T crystal) with ligation. (e) Thermally annealed motifs in a PCR tube. (f) Optical image of assembled DNA crystals on a mica surface. Scale bar: 100 µm.

### MD Simulations with OxDNA

MD computation is designed for acquiring the physical movements of particles (for example, atoms or molecules). There are two choices to balance the accuracy and simulation time. All-atom models can yield better accuracy but take longer time. Coarse-grained models use a pseudo-atom to represent a group of atoms for faster computation with less precision. Regardless of models, the particles are subjected under interactions for a given duration of time, thus generating a view of the dynamic evolution of the whole system. By numerically solving Newton’s equations of motion for the particles, their trajectories are determined.

The MD simulations based on a coarse-grained model were performed with oxDNA^33-36^ to study equilibrium conformations of the DNA crystals and their gradual structural deformations under tensile forces. Specifically, the oxDNA2 model was used with the temperature at 300 K. In typical experiments, Mg^2+^ is used to mitigate the repulsion between negative charges on DNA strands. Thus, the salt concentration was set at [Na^+^] = 0.5 M,^33,37-39^ since there is no Mg^2+^ concentration in the oxDNA simulations. We created the motif in the caDNAno2 and converted into topology and configuration files as the initial conformation in the oxDNA simulation platform with actual sequence information. The internal stresses in the single 4T motif were relaxed by MD simulations as shown in Figure 1(c). Due to the slight mismatch created by this alternating over-and-under arrangement, each direction deformed slightly. The motifs were duplicated into 125 units and combined into a 5×5×5 crystal (5 motifs in each independent direction) by both associating the hydrogen bonds (sticky-end association) and linking the covalent bonds (ligation) together at the sticky ends (Figure 1(d)). This process was realized by changing the topology and configuration files simultaneously and by monitoring the structure on an online platform, oxView.^40^ There are 25 dsDNA strands along each direction. The strands in the other two directions form 5 layers of networks which the 25 dsDNA strands path through. They serve as the cross-sectional planes. Due to the motif design, sticky-end association and ligation, there is only one strand in each dsDNA duplex that has no nicks. For example, the red, green, and blue segments at the bottom in Figure S4(a) through (c) respectively will be intact strands after ligation.

The configuration of the crystal was then relaxed with the 25 nucleotides fixed on the bottom plane (1 nucleotide per motif or per dsDNA at the outer surface of the cross-sectional plane at the bottom) immobilized using a set of force planes as reported previously^41^. The relaxation was performed for 8×10^6^ steps to obtain the initial equilibrium structure. If a native crystal was subject to relaxation, it would quickly lose hydrogen bond connections at the sticky ends. In 1×10^6^ steps, we observed motifs at the corners detached from a native crystal. Given sufficient simulation steps, the dissociation can propagate. Therefore, motifs can leave from the edges and then the outside surfaces toward the inside. Ultimately, it would disintegrate completely. Given unstable crystal conditions, we did not apply loading on the native crystals.

The relaxed ligated crystal was subjected to tensile loading by exerting forces from another set of force planes on 25 nucleotides in the top plane. The difference between the loading plane (top) and the holding plane (bottom) is the spring constant of the associated force planes. Therefore, there is one nucleotide is subject to the force from the bottom plane and one nucleotide from the top plane in each of the 25 dsDNA strands. Choosing the loading points on two different strands in the same duplex would limit the maximum possible pulling after dissociation. Thus, the two nucleotides for the top/bottom planes were chosen from the intact strand of the duplex.

The specimen in traditional tensile loading experiments is normally held at the bottom while pulled at the top. The bottom is then assumed to be stationary. We adopted the same approach in the simulations by enforcing the force constant of the holding plane to be at least 100 times that of the loading plane. In such a case, the nucleotides on the holding plane may be considered stationary along the loading direction. By moving the loading plane further from the holding plane and increasing the force constant, the crystal deformed progressively and eventually became ruptured. For any given loading plane and force constant, MD simulations were performed for 1×10^6^ steps. It may take about 3×10^5^ steps to reach the equilibrium. We defined the equilibrium as the relative fluctuations were less than 3%, and analyzed the data after reaching the equilibrium. The total running time required for each case (a combination of a loading plane position and a force constant) was about a day. After the computation, the results were analyzed by visualizing the crystal in cogli2 and by extracting the directional and positional data. The lengths of the edges in the cross-section were measured by using positional data to obtain a vector pointing from the nucleotide at one terminal of the edge to another. The angle between two edges was determined by arccosine of the dot product of the two directional vectors of the two edges, so that the cross-sectional area was acquired. Forces were recovered by product of distances from the positional data and force constants. Extension was also calculated from the positional data.

We also acquired the data of hydrogen bond energies. A threshold was set to distinguish the states of hybridization sites (bonded or broken). The value was two standard deviations higher than the averaged bond energy (a negative value) when the crystal was in the fully relaxed state (when all the bonds were supposed to be intact). The averaged bond energy and the threshold were −2.9×10^−20^ and −1.2 ×10^−20^ J, respectively. The threshold magnitude is less than a half the magnitude of the bond energy, which is reasonable. On each loaded dsDNA in the 4T crystal, there were 185 bases between the top and bottom planes. With 25 loaded duplexes, there were 4625 bases. For each result, the total bonded number was counted by comparing each hydrogen bond energy of the base with the threshold. The ratio was defined as the total bonded number divided by total base number. The 2T crystal was analyzed in a similar manner. In a 2T crystal, each loaded duplex had 91 bases and the total number of bases in loaded direction was 2275.

### Preparations of Motifs and Crystals in Experiments

In experiments, the motifs were prepared by thermal annealing of 7 strands (sequence information given in the SI) in buffered solution (Figure 1(e)). The strands were mixed with 1× TAEM buffer (an aqueous solution of 40 mM trisaminomethane or Tris, 20 mM acetic acid, 2 mM ethylenediaminetetraacetic acid or EDTA disodium salt, and 12.5 mM magnesium acetate at pH ∼8) to yield a final concentration of 2.0 µM in 40 µL. The mixture was then held for 30 min for each temperature steps of 95, 65, 50, 37, and 22°C, respectively.

The DNA crystals were assembled by hanging drop vapor diffusion method: an aliquot of 2∼5 µL assembled motif solution was incubated against 600 μL of 5× TAEM buffer in a hanging drop setup at room temperature (∼22 °C) for 3 to 7 days.^30^ For the preparation of the assembled crystals, two concepts were used: (1) ‘transferring’: the crystal was taken out by cryoloop, with minimal buffer, into another new drop; (2) ‘washing’: most of the original buffer was pipetted away from the crystals (leaving only 1∼2 µL), and the new buffer was added to the crystals. The assembled crystals were washed twice with 5× TAEM buffer and then incubated at room temperature overnight in the freshly prepared ligation mixture (an aqueous solution of 1 mM adenosine triphosphate or ATP and 80 units/μL T4 DNA ligase from New England Biolabs in 5× TAEM). After incubation, the crystals were transferred 3 times into 10 µL 5× TAEM to wash off the ligase mixture, and then they were washed twice by DI water before storage.

In the 4T motif, the center cavity is a triangle with the edge length of 17 bp or ∼5.6 nm (Figure S1). The gap between center cavities in the crystal is 25 bp or ∼8.3 nm. The edge length and the gap of the 2T motif are approximately 2.3 and 4.6 nm, respectively (Figure S2). T4 DNA ligase has a diameter of ∼5 nm if represented as a sphere.^31^ Due to the small pore size, T4 DNA ligase may not enter the 2T crystal to function. Thus, only 4T crystals with full ligation were prepared for the experiment. For the AFM measurements, the crystals need to be stable on a surface. If all three directions had the same length, the crystal (designed as a tilted cube rather than a perfect cube) could rotate upon external compression loading. Thus, the thickness (the height of the crystal) is desired to be shorter than those of the in-plane directions. Therefore, we chose the design with different sticky-end association free energies in three directions. The weakest direction is the red direction, which is the thickness and the direction to be loaded in the experiments. Typically, the crystal is around 100 µm in two in-plane directions and 10 – 20 µm in thickness as shown in Figure 1(f).

### AFM Nanoindentation Measurements

During the last transfer, the ligated 4T crystals were transferred onto freshly cleaved mica surfaces. Approximately 60 μL DI water was added for AFM measurement. AFM imaging was performed in fluid in the Peak-Force Quantitative Nano-mechanics mode with a Bruker Dimension Icon AFM and ScanAsyst-Fluid+ probes. This mode executes nanoindentation at each imaged point in such a way that the imaging and force-capturing are performed simultaneously. The Z direction (height direction) was calibrated, and the deflection sensitivity, spring constant, and tip radius were measured for accurate results. To further verify, we carried out nanoindentation measurements on several polydimethylsiloxane (PDMS) samples, and the results agreed with previous research (see SI). Visible light was shone from the bottom of the DNA crystals to the lens of visible light (rather than from the lens to the top of the DNA crystals and reflected back to the lens) for better visibility. During the AFM measurement, we made sure that the crystals were stationary and that they were always under the probe. Since the height of the crystals is about 10 – 20 μm, we adjusted the max force to 3 – 5 nN to ensure that the indentation depth was approximately 100 nm. Thus, the depth was at the level of 1% of the total height. In general, the relative indentation depth should be less than 10% with respect to the thickness, but it also must be deep enough to probe the properties of the inside part rather than just the surface. In our crystal, the maximum thickness of each layer is approximately 14 nm, and thus ∼100 nm contains at least 7 layers and is therefore a reasonable depth. For each DNA crystal, at least 7 different places were imaged with scanning scale of 2 × 2 μm for each measurement.

## RESULTS

### Externally Forced Elongation of a 4T DNA Crystal in MD Simulations

When applying tensile forces, the nucleotides on the bottom (holding) plane were immobilized while the nucleotides on the top (loading) plane were pulled away in the opposite direction. The 4T motifs were labeled from 1 to 125, and the 1 – 25 motifs were assigned on the bottom plane as illustrated in Figure 2(a). The arrow indicates the strands in three directions from 5’ to 3’ end (circle crosshair means the arrow going into the page). Similarly, the top plane has the motifs 101 – 125. The initial height was found by enforcing the holding plane for relaxation in the simulation. Once the crystal was fully relaxed on the plane, a height of ∼45 nm was observed. This is notably shorter than the height estimated based on the B-form length (approximately 61 nm based on 0.332 nm/bp). There are two major factors for the difference between the relaxed height and the B-form length. One is the curvature of the dsDNA strands. After relaxation, the strands in Figure 2(b) appear to be in a curved form compared with those in Figure 1(d). For example, two dsDNA duplexes are marked by purple arrows which have significant curvature. This is due to the nature of relaxed polymer (see the discussion below in Analysis of Force-Response). The other factor is the angle between the strands and the cross-sectional planes. The crystal is a tilted cube rather than a perfect one. The angles between the neighboring directions are all slightly over 110° in a relaxed state.^31^ Therefore, the height must be shorter than the B-form length. Based on this relaxed configuration, tensile forces were applied by implementing both loading and holding planes as indicated by a pair of black arrows.

**Figure 2.**
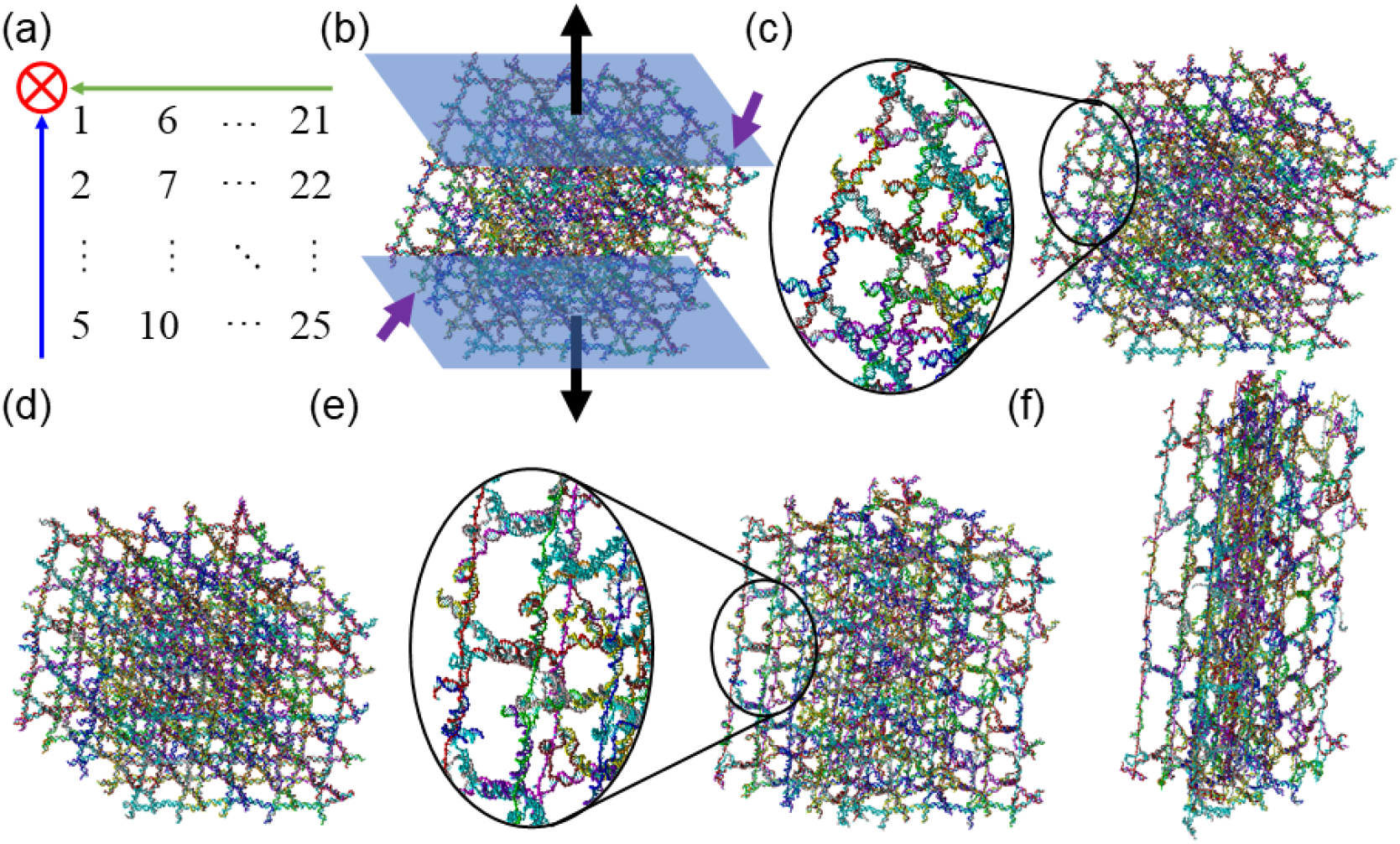
Tensile loading of a 4T DNA crystal (made of 5×5×5 4T motifs) in the MD simulations. (a) Arrangement of the motifs on the bottom holding plane. The arrows represent the strands in three directions from 5’ to 3’ end. The circle crosshair indicates the arrow going into the page. (b) The crystal in the fully relaxed state with the holding plane enforced on 25 nucleotides on the bottom. Thus, the crystal would not move perpendicular to the plane. The bottom and top planes are shown in blue. The top plane is pulled gradually, while the bottom plane is held in place. The black arrows indicate the directions of pulling forces. The purple arrows depict the examples of curved dsDNA strands. (c)-(f) The crystal under tensile loading at different stages with increased extension: (c) curved, (d) straight, (e) dsDNA dissociation, and (f) ssDNA stretch.

During the initial loading, dsDNA helices started to un-curve and straighten up (Figure 2(c)). As depicted by the insert, the strands still have curvature to some extent. This is similar to pulling a slack rope: at the very first stage, the rope deforms with little resistance. As more tensile forces were applied, the loaded dsDNA strands became nearly straight (Figure 2(d)). From this point on, the gaps between the stacked bases increased progressively upon pulling. At some point, the base-pairing started to have visible dissociation. The loaded double helices dissociated, unwound, and freed ssDNA strands. The single strands became straight as shown in Figure 2(e). This was a gradual process, which means that some of the regions were straight and dissociated while the hydrogen bonds in other regions were still intact. The distances between the 5 cross-sectional planes formed by two non-loaded directions increased noticeably. Meanwhile, the cross-sectional area shrunk. This phenomenon may be depicted by Poisson’s ratio (*ν*):

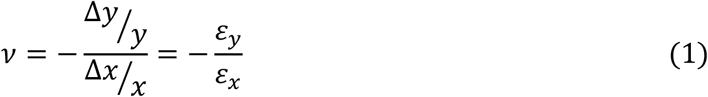

Where *ε*_*x*_ and *ε*_*y*_ are the strains in the loading (*x*) and orthogonal, non-loading (*y*) directions. The 4T crystal has positive values of the ratio, indicating the orthogonal directions contracted as the loading direction extended. Toward the end of the pulling, loaded ss-strands were almost straight but not really separated from the rest of the crystal (Figure 2(f)). The complementary strands still wrapped around them. The distances between the 5 cross-sectional planes extended significantly, and they did not detach from the loaded ssDNA completely. The connection was a few hydrogen bonds. This indicates that the crystal was loose (or able to hold significate amount of deformation) as the force provided by the few hydrogen bonds were sufficient to link the cross-sectional planes. Even the structure deformed further, it was still held together (see SI). The strands in cross-sections were severely compressed and moved out of plane. This is analogous to a flat surface that starts to pleat; it will be easier to deform further (Figure S6). This compression also confirms the positive Poisson’s ratio (Figure S8). Shortly after this point, the ssDNA broke and the simulation ended.

### Analysis of Force-Response

The applied forces and respective extensions were extracted from the simulations and plotted in Figure 3(a). There were four distinct stages with different deformation modes. In stage (i), the extension increased without any significant change in the force. The structure was initially un-tensioned and then entered the linear range after the extension of approximately 13 nm under 0.3 nN loading. In stage (ii), the extension increased linearly with the applied force, until reaching the yield strength of ∼1.7 nN. Then, as the force increased moderately, the structure was extended drastically in stage (iii). For example, the crystal was extended for approximately 49 nm while the force was varied from 1.7 to 3.7 nN (with a slope of ∼0.04 N/m). In comparison, the linear elasticity in stage (ii) ranged from 0.27 to 1.7 nN for an extension of only ∼9 nm (with a slope of ∼0.16 N/m). When the crystal was pulled further into stage (iv), the force increased drastically until the crystal eventually broke. Representative images of the deformation modes shown in Figure 2(b)-(f) are denoted as filled black circles in Figure 3(a). All simulated structures corresponding to the plotted data are presented in Figure S6.

**Figure 3.**
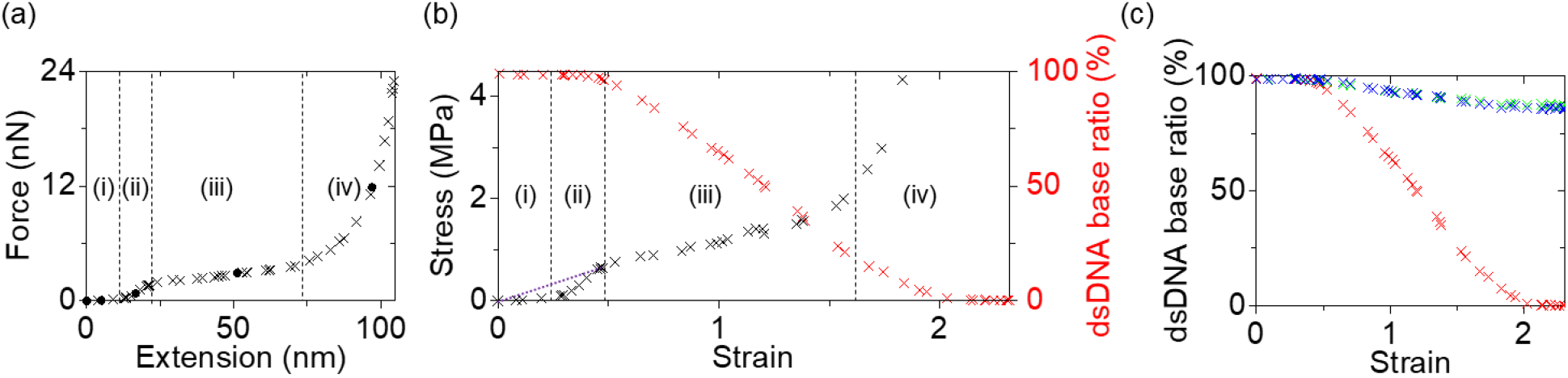
MD simulation results of a 4T crystal. (a) Force-extension plot. The cross marks all data points (presented in Figure S6), while the solid dot indicates the result corresponding to the configuration presented in Figure 2(b)-(f). (b) Stress-strain plot processed from the force-extension curve. The stress data go up to ∼22 MPa but are plotted below 4.5 MPa for clarity. The base-pairing ratio of the 25 dsDNA strands in the pulling direction is also plotted in red. The purple dotted line indicates the Young’s modulus of the structure. (a)-(b) Four deformation modes are observed as indicated by the vertical dotted lines: (i) un-tension, (ii) linear elasticity, (iii) dsDNA dissociation, and (iv) ssDNA stretch. (c) Association ratios of the dsDNA bases in all three directions. The color codes on red, green, and blue are identical in Figure 1. Note that the red indicates the pulling direction as in (b).

In mechanics, the stress-strain plot is widely used to determine physical properties of the specimen such as the Young’s modulus. Stress is defined as applied force per unit cross-sectional area. It should be noted that the cross-sectional area varied during the extension and the real time area was used for all the data points. As shown in equation 1, the strain is extension per unit reference length (which is the initial height here), thus it is proportional to extension. The stages (i) through (iv) in Figure 3(b) correspond to those in Figure 3(a). The association ratio of the dsDNA bases in the loaded 25 double helices can provide insight into the mechanisms involved in the different deformation modes. The dsDNA bases were nearly intact in stage (i). In stage (ii), the dsDNA ratio decreased slightly, yet it was greater than 96%, indicating that there were negligible dissociations. When the crystal was pulled further into stage (iii), the dsDNA ratio decreased monotonically. At the end of stage (iii), the dsDNA ratio was below 22%, indicating that most of the hydrogen bonds along the loading direction were broken. Thus, the force response in stage (iv) becomes related to ssDNA stretch rather than dsDNA. As the dsDNA strands were all dissociated, the stress and the force required for extension increased rapidly. The loaded ssDNA broke after the dsDNA ratio dropped under 1%. We termed the four stages, un-tension, linear elasticity, dsDNA dissociation, and ssDNA stretch, to represent the major deformation modes in the stages.

The association ratios of the dsDNA bases in the other two directions, green and blue, are plotted in Figure 3(c) to compare with the loading direction, red. As discussed above, the 4T motif has different sequences in three directions, and thus, the 4T crystal can be thin in one direction. However, the edge length and the strand arrangement in all three directions are the same if we neglect the nick on the cyan strand (see Figures S1 and S4). When analyzing the association ratio of the red direction, we considered the bases between the bottom and the top cross-sectional planes (Figure S5(a)). The bases protruding out of the two planes were not considered in the estimation. Similarly, the bases between the left and right cross-sections (Figure S5(b)) were taken into account for the base-pairing ratio in the green direction. For the blue direction, the bases examined were between the front and back cross-sections (Figure S5(c)). When the loading direction was almost intact in stages (i) and (ii), the other two directions were also intact. As the bases in red dissociated almost linearly in stage (iii), those in green and blue dissociated as well. However, the base-pairing ratios in the non-loaded directions were always above 85%, even in stage (iv). This shows that the non-loading directions were not severely affected. The crystal is similar to 25 loaded double helices in parallel with some effect from perpendicular directions.

### Nanoindentation Experiments on the 4T DNA Crystals

We deposited the assembled 4T DNA crystals on mica surfaces and performed AFM nanoindentation in fluid to verify the simulation results. The measured crystals were placed under the AFM probe and engaged for scanning as depicted in Figure 4(a). We acquired the surface topography (Figure 4(b)), while performing nanoindentation. There were repeating hollows on the surface, corresponding to the center pores of the connected motifs. The upper surface had some variations in height, which was within 20 nm, mostly less than 8 nm. If the indentation depth was at 100 nm level, these variations should not affect the measurement results significantly. At each imaged point, approach and retract force curves were collected and analyzed by the AFM software. Figure 4(c) presents a pair of representative force curves at a single point. The separation is defined as the displacement from the lowest indented point. As the probe approached the surface from high up, there was barely any force at the beginning. The contact point was judged by the start of monotonic increase of the force with respect to the separation. At around 100 nm of separation, the force started to rise, indicating the initial contact point of the AFM tip. Therefore, the indentation depth was approximately 100 nm. As the probe tip pressed the surface continuously, the force increased drastically, reaching the maximum force of about 3 nN. During the retraction, the probe was slowly raised up from the indented spot. The retract curve was similar to the approach process, but it went blow 0 (attraction force), indicating that the surface was sticky.

**Figure 4.**
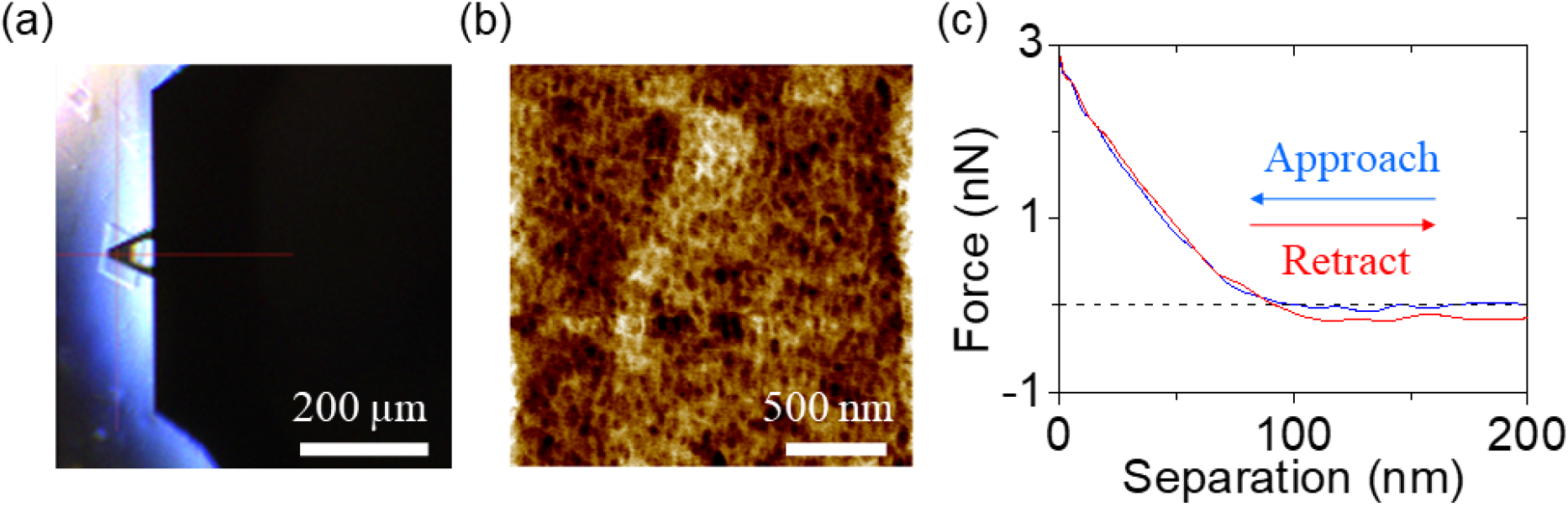
AFM nanoindentation of the 4T DNA crystal. (a) The AFM probe hovers on a DNA crystal before measurement. (b) Representative topography of the upper surface of a DNA crystal. The height variation is within 20 nm, mostly less than 8 nm. (c) A pair of representative approach and retract force curves at a single point. The separation is the displacement from the lowest indented point. The indentation depth was approximately 100 nm.

We used a conical tip model to analyze the force curves (see SI).^42^ The analysis suggests that the measured structural Young’s modulus has an average of approximately 1 MPa. This is consistent with the value from the MD simulations indicated by the inclined purple dotted line in Figure 3(b). The linear elasticity in the simulations ranges from 0.4 to 1.4 MPa. The agreement between experiments and simulations suggests that we reached the linear elasticity as the AFM tip probed into the crystal. The un-tension stage did not prevent the measurement. Limited by the maximum possible indentation depth, we could not precede further.

### Computational Studies of a 2T Crystal

We constructed another crystal from 5×5×5 2T motifs^26^ (Figure 5(a)) to investigate the effect of motif length. Similarly, the bottom holding plane was enforced to obtain the initial equilibrium conformation before loading. The corresponding height was ∼29 nm (Figure 5(b)). It is interesting to note that this height is nearly the same as the one estimated based on the B-form dsDNA (∼30 nm). Due to a higher crossover density, the dsDNA strands in this 2T crystal did not show any noticeable curvature when compared with the 4T crystal. This agrees with the height comparison. Tensile forces were applied on this relaxed crystal. Notably, there was no un-tension period. The gaps between the stacked bases increased upon loading (Figure 5(c)). With continued extension, the base-pairs started to have visible dissociation as depicted in Figure 5(d). The distance between neighboring cross-sectional planes increased significantly, while the cross-sectional area remained roughly the same. The green and blue directions were nearly independent from the loading direction. Toward the end of the pulling, the loaded ss-strands were almost separated from the rest of the crystal (Figure 5(e)). Instead of being further pulled apart from each other, the 5 cross-sectional planes failed from the 25 loaded ssDNA and stacked towards the bottom plane. There were still some base pairings that held related strands together against a full detachment. As the structure deformed further, the cross-sectional planes collapsed to the bottom (see SI). The connections between the planes were tighter than the linkages to the loaded strands at this point. After this collapsing point, the ssDNA did not stretch much more and broke, which marked the end of the simulation (Figure S7).

**Figure 5.**
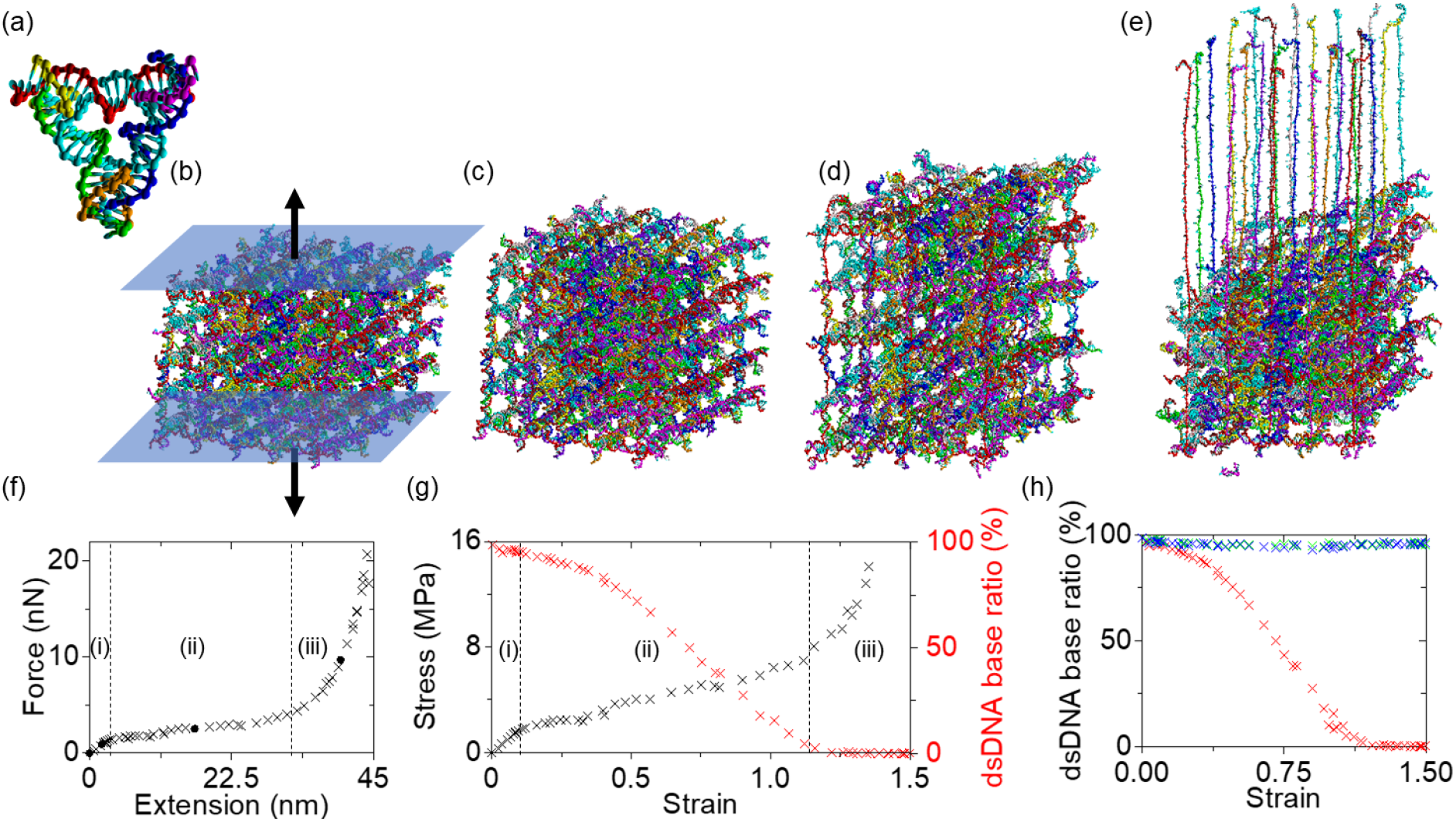
(a) Molecular model of a 2T tensegrity triangle motif in oxDNA. The color codes are the same as in Figures 1 and 2. (b) A 2T crystal (5×5×5 motifs) with ligation after full relaxation, while enforcing just the holding (bottom) plane. The top blue plane is pulled gradually, while the bottom plane (also in blue) is held in place. (c)-(e) Snapshot images of the crystal under tensile loading at different stages. (f) Force-extension plot. (g) Stress-strain curve with the base-pairing ratio of the dsDNA in the loading direction. The stress data go up to ∼30 MPa but are plotted below 16 MPa for clarity. (f)-(g) The vertical dotted lines divide the plots into 3 stages: (i) linear elasticity, (ii) dsDNA dissociation, and (iii) ssDNA stretch. The cross marks all data points (presented in Figure S7), while the solid dot indicate the configuration images in (b)-(e). (h) Base association ratio of the dsDNA in all three directions.

The force-extension, stress-strain, and base-pairing ratio were extracted from the simulations and plotted in Figure 5(f)-(h). There are three stages with distinct deformation modes: (i) linear elasticity, (ii) dsDNA dissociation, and (iii) ssDNA stretch. The vanishing of the un-tension stage is because the 2T crystal was in tension from the beginning. The structural Young’s modulus was approximately 17.6 MPa which is more than an order of magnitude larger than that of the 4T crystal. This is attributed to the elimination of the un-tension stage. The longer the un-tension stage, the lower the Young’s modulus. The yield strength was approximately 1.4 nN, which is similar to that of the 4T crystal, since they both have 25 duplexes along the loading direction. The slopes for the linear elasticity and dissociation stages (spring constants) were about 0.43 and 0.10 N/m, respectively, which are several folds greater than those of the 4T crystal. There are two reasons for this. First, the motif edge length of the 2T crystal is a half of that of the 4T crystal. Under the same loading, shorter dsDNA will be extended shorter. Second, a higher crossover density makes a tighter structure, thus the 2T crystal is harder to pull. As the crystal was extended into stage (iii), the force increased drastically until the crystal eventually broke.

The base-pairing ratio of the dsDNA strands in the loaded direction provides insights on the stress-strain behavior. In stage (i), the dissociations are negligible with the association ratio above 95%. Thus, the structural deformation is by almost pure dsDNA stretch. Then, the ratio decreases monotonically reaching below 5% at the end of stage (ii). This trend corresponds to the drastic extension of the crystal height. The association ratio at the end of the dissociation stage is much lower than 4T crystal. When the force and stress started to increase drastically with respect to extension or strain, the deformation mode shifted from dsDNA dissociation to ssDNA stretch. Ideally, the association ratio should drop to 0 by the end of the dissociation stage. The 2T crystal reached < 5%, which is a good match, indicating that the non-loaded directions do not really affect the loading direction (also see the discussion below on Figure 5(h)). In contrast, the 4T crystal reached < 22 %. This could be due to the mutual influences between the loading and non-loaded directions. With the loaded strands mostly dissociated in stage (iii), the ssDNA strands on the top pulling plane were stretched completely as shown in Figure 5(e).

Figure 5(h) depicts the association ratio of the dsDNA bases in all three directions. The non-loading directions were always above 93%. Interestingly, after the loading direction dropped below 10%, both green and blue regained some association. With the loaded ssDNA detached from the rest, the green and blue had freedom to re-hybridize with the previously dissociated complementary nucleotides. The more bases in red detached from the rest of the structure, the more bases in green and blue reassociated with the structure. This shows that the green and blue directions were less affected compared with the situation in the 4T crystal. In a tighter structure with a higher crossover density, each direction behaves more independently.

## DISCUSSION

### Experiment and Simulation

We have studied the mechanical deformation behaviors of ligated DNA crystals with MD simulations and AFM nanoindentation which together provided invaluable insights on the DNA mechanics. To the best of our knowledge, this study presents the first experimental measurement of direct force-response on complex DNA architectures. We chose indentation (compressive loading) in the experiment as it does not require extra immobilization methods for the crystals. The only challenge is the possible rotation of the crystal. It was overcome by building flat, thin crystals. As a comparison, tensile loadings would require solid attachment between the mica and the crystals. Since the loading were to be exerted by the AFM probe, DNA strands would also need to be attached to the probe for a connection with the crystal. This is similar to an extension of a single polysaccharide chain by AFM.^43^ All these steps will add a lot of complexity to the system. During the indentation experiments, we did not observe a sharp transition between the un-tension and linear elasticity, as in the simulations. If any, it would be too weak to discover under the thermal fluctuation. There are some limitations such as the maximum indentation depth and the smallest scale we can probe. Due to the limited indentation depth, plastic deformation was not achieved.

Coarse-grained MD simulations can be versatile on exploring different loading methods on various dimensions of DNA structures. We tried both tensile and compressive loadings. The immobilization for extension is simple in the simulation, but the compression can be complex. Under compressive loading, the tilted cubic crystal may rotate, as in the experiments. Building a flat, thin crystal will significantly add the number of motifs. This will require the computation power greatly. Besides, it can apply loading on the ligated 2T crystal, which was unattainable in the experiments. The simulations provided a lot of detailed information, which was not possible in the measurements. A good example is the association ratios of the dsDNA bases. Given the huge number of bases and the transient loading conditions, it is impossible to acquire the real time association ratios experimentally. Plus, the computation revealed the distinct stages during the loading until the failing of the covalent bonds. Limitations are present for simulations as well. The maximum size to be calculated is limited by the model and the computation power. A larger structure (e.g., larger than the crystal model made of 125 motifs) might provide more details and match the experimental scenarios better, but the balance between the calculation speed and insights is critical.

### 4T and 2T Crystals

Both crystals have similar motif designs and assembly methods. As a result, they are both in a titled cubic shape. Due to the motif size, the ligation of the 2T crystal was not possible in the experiment. The native 4T crystal in simulation was unstable as discussed above. The detachment of the motifs at the corners was observed early on during the initial relaxation (within first 10^6^ simulation steps). The crossover density in the 2T crystal is twice as that in the 4T crystal. Therefore, it is a tighter crystal, there will be more internal stresses, and we expect more severe dissociation in the native 2T crystal. We found that approximately 1/3 of the sticky-end connections between the bottom cross-section and its neighboring plane came off during the initial relaxation. Similar dissociation pattern was observed with the top cross-sectional planes. This is consistent with experimental observation that both crystals disintegrate under normal salt concentrations and high ion strength (*e.g*., 50 mM Mg^2+^) is needed for stabilizing the native crystals.

Despite many similarities, the 4T and 2T crystals show interesting differences in mechanical properties. As discussed above, the structural Young’s moduli of the 4T and 2T crystals are approximately 1 and 17.6 MPa, respectively. This difference is mostly caused by the elimination of the un-tension stage in the 2T crystal. In terms of the yield force, however, they are similar; about 1.7 and 1.4 nN for the 4T and 2T crystals. This is because the yield strength is simply the total force exerted on the 25 double helices. The linear elasticity in the 2T crystal starts without the un-tension, whereas the 4T crystal starts at ∼0.3 nN. For a fair comparison, we may exclude the force related to the un-tension stage. Then, both crystals have the yield force of 1.4 nN.

### Mechanical Behaviors of a Structure and its Components

Figure 6 compares the deformation behaviors of a single dsDNA and averaged dsDNA strands in the 2T and 4T DNA crystals. A general trend of pulling a ds-λ-DNA in Figure 6(a) was obtained from previous reports in the literature. Caron and coworkers attached one end of the λ-DNA on a latex bead subjected to a micropipette for extension of the strand.^44^ The other terminal was connected to a fiber holder. The force was measured by the fiber holder while the displacement was acquired by the controlled microbead. By pulling the bead away from the holder, the λ-DNA was extended. Bustamante et al. linked two beads to the either ends of a λ-DNA strand.^45^ One was immobilized by a pipette tip while the other was subjected to an optical tweezer. The force and the extension were both recovered from the optical inputs. There are also other methods for probing DNA mechanics including hydrodynamic drag, magnetic beads, and glass needles, as summarized by Bustamante et al.^46^ and Cocco and coworker^47^. Regardless of the methods, the results of λ-DNA extension are comparable. Four regimes were observed: entropic elasticity, intrinsic elasticity, overstretching transition, and breaking covalent bonds.

**Figure 6.**
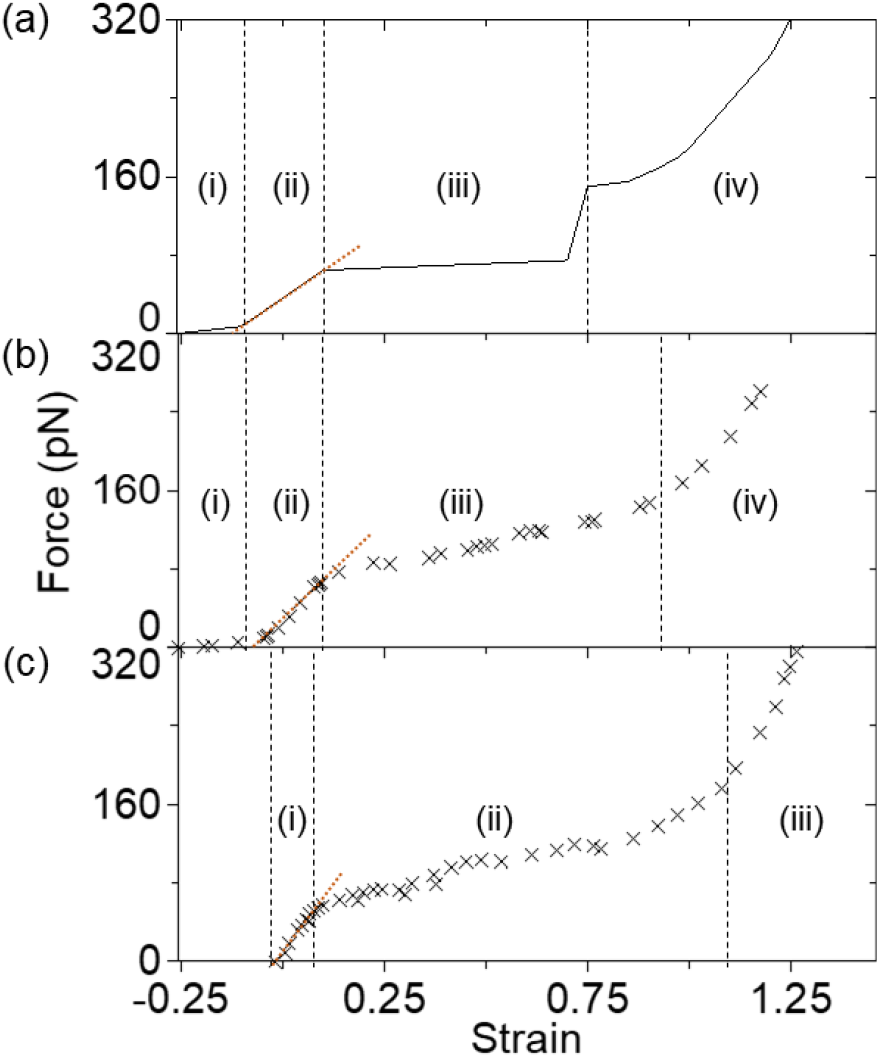
(a) General trend of pulling a ds-λ-DNA from previous publications^44-47^. The strain data start at −1 but are plotted from −0.25 for clarity. Before the linear elastic range, the DNA molecule is very soft at the beginning (regime (i) or entropic elasticity), and it enters the intrinsic elasticity at ∼8 pN. The extension on the dsDNA backbone continues until ∼65 pN (regime (ii)). Beyond this point, the overstretching transition (regime (iii)) is followed by a sharp increase up to ∼150 pN at the end. In regime (iv) (termed breaking covalent bonds), the force increases exponentially. The maximum force one dsDNA could hold is approximately 1 nN. (b)-(c) Force-strain plots of averaged dsDNA strands in the 4T and 2T crystals. The plotted force is the pulling force per loaded dsDNA. Thus, they are the forces in Figure 3(a) and Figure 5(f) divided by 25, respectively. The force data go up to ∼1 nN but are plotted below 320 pN for clarity. The strains are re-defined by the reference length of B-form DNA, for the comparison with the literature. Therefore, the initial strain is less than 0 since the relaxed length is shorter than the B-form length. The maximum forces per dsDNA are approximately 0.9 and 0.8 nN (not shown in (b) and (c)). The brown dotted lines in (a)-(c) indicate the slope of the linear elasticity.

If we compare the force-response of a single dsDNA with the averaged dsDNA in the DNA crystal (Figure 6(b)-(c)), the overall magnitudes of the transition points between the regimes and stages share some similarities. The deformation mechanism for each regime corresponds to that of the stage at similar strain. However, they are not completely identical. In the entropic elasticity of a single duplex, it starts from the strain of almost −1 since it has no restriction against tangling together. In contrast, the dsDNA strand in the crystal is restricted by other helices, especially the helices in different directions. The 4T crystal starts the un-tension stage from the strain of −0.25, and the 2T crystal starts at 0. In the overstretching transition regime, the single dsDNA has a drastic change of force in the end, which is not reflected by the averaged dsDNA of the crystal in the dissociation stage. There are variations on the force distribution over the cross-sectional areas of the crystal. This leads to a steady increment rather than a drastic change of the force-reaction. Moreover, part of the impact may be absorbed with the non-loading directions. With the understanding of the difference between Figure 6(a) and (b)-(c), we conclude that dsDNA is one of the determinators of the properties of DNA crystals. By assembling dsDNA into a crystal, the force-reaction curve is smoother.

Another difference between the structures and components is the Young’s moduli. The slope in the intrinsic elasticity (noted by the brown dotted line in Figure 6(a)) of a single dsDNA is about 290 pN. The slopes per strand in the 4T and 2T crystals are approximately 550 and 390 pN, respectively. Given the cross-section of a dsDNA strand of roughly 3.8 nm^2^, the Young’s moduli from the linear elasticity in Figure 6(a)-(c) are approximately 76, 145 and 102 MPa. This is consistent with typical Young’s modulus (on the order of 10^2^ MPa) of dsDNA reported in the literature.^48^ These values are one to two orders of magnitude higher than those of the crystals (1 and 17.6 MPa). Note that the Young’s moduli of the crystals are the structural properties and are defined as force per area per strain, that is, the slope in the force-strain plot divided by the cross-sectional area. The cross-section of a DNA crystal has the area that includes both components (dsDNA) and cavity (where there are no dsDNA). Overall, the comparison clearly shows the intricate correlation between a structure and its components (dsDNA rods) in mechanical properties. However, the deformation behaviors can be inferred only partially, and the architecture must be considered along with motif designs.

### Outlook

We found that DNA crystals made of tighter and shorter motifs will have a higher crossover density and likely will not have the un-tension stage. By the same token, we can infer that a 3T crystal with ligation will have some un-tension region but less than that of the 4T crystal. By varying the crossover density, it may be possible to assemble a crystal with motifs of different lengths (e.g., 2T and 4T combined). Such a heterogenous crystal could offer interesting properties with some parts tight and stiff, while other parts more relaxed un-tensioned (thereby more deformation tolerant).

The plastic deformation of DNA crystals is initiated by the dissociation of hydrogen bonds. There are other ways to induce dissociation; for example, heating or adding urea or other similar chemicals. It is thus possible to study the transition from elastic to plastic deformations by controlled dissociation as a function of urea concentration (although dissociation will not be directional). More dissociation means more extension if the loading is kept constant. Similarly, DNA crystals could be used as force-based temperature sensors. They will deform visibly once the temperature passes a certain point if the structure is pre-loaded with certain force. The temperature point may be altered by adjusting the preload.

For a ligated DNA crystal, the total failing of the structure occurs when the covalent bonds on the backbone are pulled apart. In such a case, the breaking point may be random with perfect ligation. The failing points may be introduced at certain locations. Introducing imperfect ligation on the crystal (that is, leaving some sticky-ends in native form) would be useful. The ligated part can be solid and should be able to withhold mechanical forces, while the native segment may collapse upon external loads. Further, it is possible to achieve patterned ligation on a DNA crystal. We recently demonstrated complex yolk-shell and Matryoshka-doll shaped ligation patterns within DNA crystals by controlling the ligation process.^31^ DNA crystals with such patterned ligation could have directional mechanical properties and serve as a platform toward programmable actuation devices.

## ACKNOWLEDGEMENT

This work was supported by the U.S. National Science Foundation (NSF) under the award no. 2025187 (4T DNA crystals) and by the U.S. Department of Energy (DOE), Office of Science, Basic Energy Sciences (BES) under award no. DE-SC0020673 (2T crystals).

## Notes

### Competing Interest Statement

The authors have declared no competing interest.

## REFERENCES

1 Seeman, N. C. & Sleiman, H. F. DNA Nanotechnology. Nature Reviews Materials 3, 1–23 (2017).

2 Pinheiro, A. V., Han, D., Shih, W. M. & Yan, H. Challenges and Opportunities for Structural DNA Nanotechnology. Nature Nanotechnology 6, 763–772 (2011).

3 Seeman, N. C. Nucleic Acid Junctions and Lattices. Journal of Theoretical Biology 99, 237–247 (1982).

4 Lin, C., Liu, Y., Rinker, S. & Yan, H. DNA Tile Based Self-Assembly: Building Complex Nanoarchitectures. ChemPhysChem 7, 1641–1647 (2006).

5 Park, S. H., Pistol, C., Ahn, S. J., Reif, J. H., Lebeck, A. R., Dwyer, C. & LaBean, T. H. Finite-Size, Fully Addressable DNA Tile Lattices Formed by Hierarchical Assembly Procedures. Angewandte Chemie 118, 749–753 (2006).

6 Choi, J., Chen, H., Li, F., Yang, L., Kim, S. S., Naik, R. R., Ye, P. D. & Choi, J. H. Nanomanufacturing of 2D Transition Metal Dichalcogenide Materials Using Self-Assembled DNA Nanotubes. Small 11, 5520–5527 (2015).

7 Rothemund, P. W. K. Folding DNA to Create Nanoscale Shapes and Patterns. Nature 440, 297–302 (2006).

8 Powell, J. T., Akhuetie-Oni, B. O., Zhang, Z. & Lin, C. DNA Origami Rotaxanes: Tailored Synthesis and Controlled Structure Switching. Angewandte Chemie International Edition 55, 11412–11416 (2016).

9 Wei, B., Dai, M. & Yin, P. Complex Shapes Self-Assembled from Single-Stranded DNA Tiles. Nature 485, 623–626 (2012).

10 Ke, Y., Ong, L. L., Shih, W. M. & Yin, P. Three-Dimensional Structures Self-Assembled from DNA Bricks. science 338, 1177–1183 (2012).

11 Ke, Y., Ong, L. L., Sun, W., Song, J., Dong, M., Shih, W. M. & Yin, P. DNA Brick Crystals with Prescribed Depths. Nature Chemistry 6, 994–1002 (2014).

12 Lin, Y., Shen, X., Wang, J., Bao, L., Zhang, Z. & Pang, D. Measuring Radial Young’s Modulus of DNA by Tapping Mode AFM. Chinese Science Bulletin 52, 3189–3192 (2007).

13 Nguyen, T.-H., Lee, S.-M., Na, K., Yang, S., Kim, J. & Yoon, E.-S. An Improved Measurement of DsDNA Elasticity Using AFM. Nanotechnology 21, 075101 (2010).

14 Li, L., Liu, L., Tabata, O. & Li, W. J. in The 9th IEEE International Conference on Nano/Micro Engineered and Molecular Systems (NEMS). 684–687 (IEEE).

15 Ma, Z., Kim, Y.-J., Park, S., Hirai, Y., Tsuchiya, T., Kim, D.-N. & Tabata, O. in 10th IEEE International Conference on Nano/Micro Engineered and Molecular Systems. 581–584 (IEEE).

16 Zhou, F., Sun, W., Zhang, C., Shen, J., Yin, P. & Liu, H. 3D Freestanding DNA Nanostructure Hybrid as a Low-Density High-Strength Material. ACS Nano 14, 6582–6588 (2020).

17 Goodman, R. P., Schaap, I. A., Tardin, C. F., Erben, C. M., Berry, R. M., Schmidt, C. F. & Turberfield, A. J. Rapid Chiral Assembly of Rigid DNA Building Blocks for Molecular Nanofabrication. Science 310, 1661–1665 (2005).

18 Li, R., Chen, H. & Choi, J. H. Topological Assembly of a Deployable Hoberman Flight Ring from DNA. Small 17, 2007069 (2021).

19 Li, R., Chen, H. & Choi, J. H. Auxetic Two-Dimensional Nanostructures from DNA. Angewandte Chemie 60, 7165–7173 (2021).

20 Zhang, Z., Yang, Y., Pincet, F., Llaguno, M. C. & Lin, C. Placing and Shaping Liposomes with Reconfigurable DNA Nanocages. Nature Chemistry 9, 653–659 (2017).

21 Powell, J. T., Akhuetie-Oni, B. O., Zhang, Z. & Lin, C. DNA Origami Rotaxanes: Tailored Synthesis and Controlled Structure Switching. Angewandte Chemie 128, 11584–11588 (2016).

22 Chen, H., Zhang, H., Pan, J., Cha, T.-G., Li, S., Andréasson, J. & Choi, J. H. Dynamic and Progressive Control of DNA Origami Conformation by Modulating DNA Helicity with Chemical Adducts. ACS Nano 10, 4989–4996 (2016).

23 Ke, Y., Bellot, G., Voigt, N. V., Fradkov, E. & Shih, W. M. Two Design Strategies for Enhancement of Multilayer–DNA-Origami Folding: Underwinding for Specific Intercalator Rescue and Staple-Break Positioning. Chemical Science 3, 2587–2597 (2012).

24 Zhao, Y.-X., Shaw, A., Zeng, X., Benson, E., Nyström, A. M. & Högberg, B. DNA Origami Delivery System for Cancer Therapy with Tunable Release Properties. ACS nano 6, 8684–8691 (2012).

25 Miller, H. L., Contera, S., Wollman, A. J., Hirst, A., Dunn, K. E., Schröter, S., O’Connell, D. & Leake, M. C. Biophysical Characterisation of DNA Origami Nanostructures Reveals Inaccessibility to Intercalation Binding Sites. Nanotechnology 31, 235605 (2020).

26 Zheng, J., Birktoft, J. J., Chen, Y., Wang, T., Sha, R., Constantinou, P. E., Ginell, S. L., Mao, C. & Seeman, N. C. From Molecular to Macroscopic via the Rational Design of a Self-Assembled 3D DNA Crystal. Nature 461, 74–77 (2009).

27 Simmons, C. R., Zhang, F., Birktoft, J. J., Qi, X., Han, D., Liu, Y., Sha, R., Abdallah, H. O., Hernandez, C., Ohayon, Y. P., Seeman, N. C. & Yan, H. Construction and Structure Determination of a Three-Dimensional DNA Crystal. Journal of the American Chemical Society 138, 10047–10054 (2016).

28 Zhang, F., Simmons, C. R., Gates, J., Liu, Y. & Yan, H. Self-Assembly of a 3D DNA Crystal Structure with Rationally Designed Six-Fold Symmetry. Angewandte Chemie International Edition 57, 12504–12507 (2018).

29 Nguyen, N., Birktoft, J. J., Sha, R., Wang, T., Zheng, J., Constantinou, P. E., Ginell, S. L., Chen, Y., Mao, C. & Seeman, N. C. The Absence of Tertiary Interactions in a Self-Assembled DNA Crystal Structure. Journal of Molecular Recognition 25, 234–237 (2012).

30 Li, Z., Zheng, M., Liu, L., Seeman, N. C. & Mao, C. 5’-Phosphorylation Strengthens Sticky-End Cohesions. Journal of the American Chemical Society 143, 14987–14991 (2021).

31 Li, Z., Liu, L., Zheng, M., Zhao, J., Seeman, N. C. & Mao, C. Making Engineered 3D DNA Crystals Robust. Journal of the American Chemical Society 141, 15850–15855 (2019).

32 Zhao, J., Chandrasekaran, A. R., Li, Q., Li, X., Sha, R., Seeman, N. C. & Mao, C. Post-Assembly Stabilization of Rationally Designed DNA Crystals. Angewandte Chemie International Edition 54, 9936–9939 (2015).

33 Snodin, B. E. K., Randisi, F., Mosayebi, M., Šulc, P., Schreck, J. S., Romano, F., Ouldridge, T. E., Tsukanov, R., Nir, E., Louis, A. A. & Doye, J. P. K. Introducing Improved Structural Properties and Salt Dependence into a Coarse-Grained Model of DNA. Journal of Chemical Physics 142, 234901 (2015).

34 Šulc, P., Romano, F., Ouldridge, T. E., Rovigatti, L., Doye, J. P. & Louis, A. A. Sequence-Dependent Thermodynamics of a Coarse-Grained DNA Model. The Journal of Chemical Physics 137, 135101 (2012).

35 Rovigatti, L., Šulc, P., Reguly, I. Z. & Romano, F. A Comparison Between Parallelization Approaches in Molecular Dynamics Simulations on GPUs. Journal of Computational Chemistry 36, 1–8 (2015).

36 Ouldridge, T. E., Louis, A. A. & Doye, J. P. Structural, Mechanical, and Thermodynamic Properties of a Coarse-Grained DNA Model. The Journal of Chemical Physics 134, 02B627 (2011).

37 Snodin, B. E., Romano, F., Rovigatti, L., Ouldridge, T. E., Louis, A. A. & Doye, J. P. Direct Simulation of the Self-Assembly of a Small DNA Origami. ACS Nano 10, 1724–1737 (2016).

38 Engel, M. C., Smith, D. M., Jobst, M. A., Sajfutdinow, M., Liedl, T., Romano, F., Rovigatti, L., Louis, A. A. & Doye, J. P. K. Force-Induced Unravelling of DNA Origami. ACS Nano 12, 6734–6747 (2018).

39 Snodin, B. E., Schreck, J. S., Romano, F., Louis, A. A. & Doye, J. P. Coarse-Grained Modelling of the Structural Properties of DNA Origami. Nucleic Acids Research 47, 1585–1597 (2019).

40 Poppleton, E., Bohlin, J., Matthies, M., Sharma, S., Zhang, F. & Šulc, P. Design, Optimization and Analysis of Large DNA and RNA Nanostructures through Interactive Visualization, Editing and Molecular Simulation. Nucleic Acids Research 48, e72 (2020).

41 Li, R., Chen, H., Lee, H. & Choi, J. H. Elucidating the Mechanical Energy for Cyclization of a DNA Origami Tile. Applied Sciences 11, 2357 (2021).

42 Hainsworth, S., Chandler, H. & Page, T. Analysis of Nanoindentation Load-Displacement Loading Curves. Journal of Materials Research 11, 1987–1995 (1996).

43 Grandbois, M., Beyer, M., Rief, M., Clausen-Schaumann, H. & Gaub, H. E. How strong is a covalent bond? Science 283, 1727–1730 (1999).

44 Cluzel, P., Lebrun, A., Heller, C., Lavery, R., Viovy, J.-L., Chatenay, D. & Caron, F. DNA: an Extensible Molecule. Science 271, 792–794 (1996).

45 Smith, S. B., Cui, Y. & Bustamante, C. Overstretching B-DNA: The Elastic Response of Individual Double-Stranded and Single-Stranded DNA Molecules. Science 271, 795–799 (1996).

46 Bustamante, C., Smith, S. B., Liphardt, J. & Smith, D. Single-Molecule Studies of DNA Mechanics. Current Opinion in Structural Biology 10, 279–285 (2000).

47 Marko, J. F. & Cocco, S. The Micromechanics of DNA. Physics World 16, 37 (2003).

48 Hogan, M. & Austin, R. H. Importance of DNA Stiffness in Protein–DNA Binding Specificity. Nature 329, 263–266 (1987).

