## Supplement for "Mechanical Deformation Behaviors and Structural Properties of Ligated DNA Crystals"

**Content**

S1. DNA Sequence Information
S2. Crystal Assemblies and Arrangements
S3. Supporting Simulation Snapshots
S4. Supporting Processed Simulation Data
S5. Nanoindentation Concepts and Verifications
S6. References

### S1. DNA Sequence Information

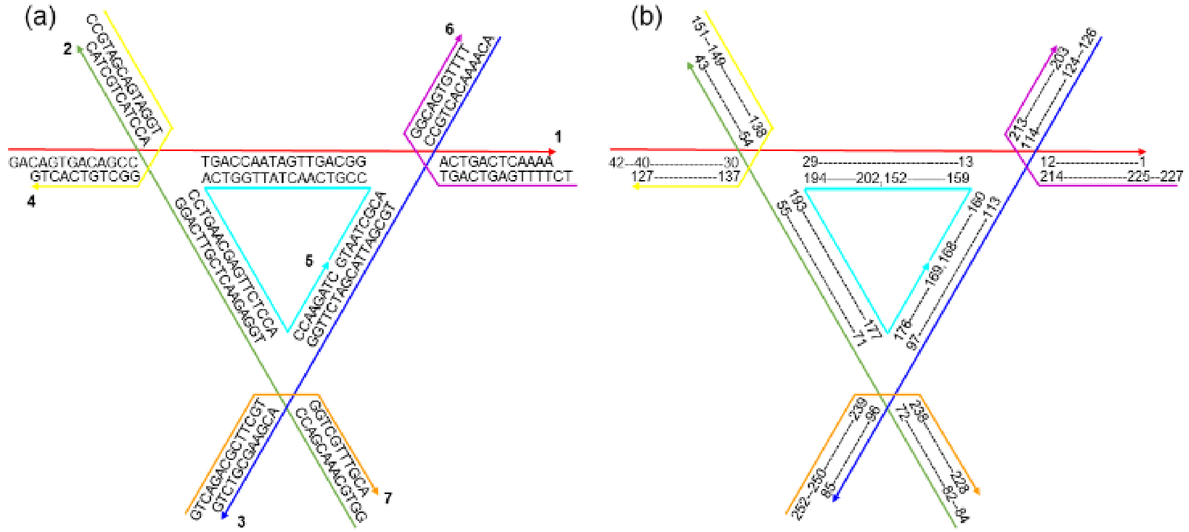


**Figure S1.** Schematics of the 4T motif. The color scheme is the same as in Figure 1. (a) The sequence information. The strands are labelled from 1 to 7, all at the 3' end. (b) The labelling of each nucleotide (nt) in the oxDNA simulation, all from 3' to 5' end. There are 252 nt in total.

The sequences of strands are as listed, all from 5' to 3' end:
1: GACAGTGACAGCCTGACCAATAGTTGACGGACTGACTCAAAA
2: GGTGCAAACGACCTGGAGAACTCGTTCAGGACCTACTGCTAC
3: ACAAAACACTGCCTGCGATTACGATCTTGGACGAAGCGTCTG
4: CCGTAGCAGTAGGTGGCTGTCACTG
5: GTAATCGCACCGTCAACTATTGGTCACCTGAACGAGTTCTCCACCAAGATC
6: TCTTTTGAGTCAGTGGCAGTGTTTT
7: GTCAGACGCTTCGTGGTCGTTTGCA

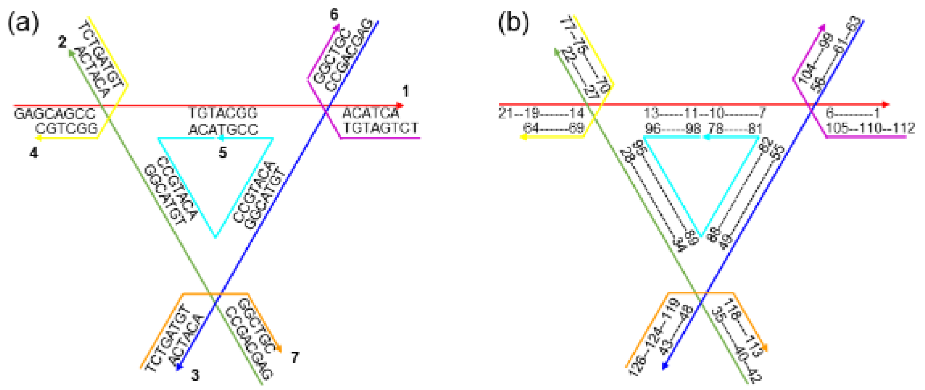


**Figure S2.** Schematics of the 2T motif. The color scheme is the same as in Figure 5. The design and strand arrangement are similar to those in Figure S1, but shorter strands are used (126 nt in total).

The sequence information (from 5' to 3' end):
1, 2, and 3: GAGCAGCCTGTACGGACATCA
4, 6, and 7: TCTGATGTGGCTGC
5: ACACCGTACACCGTACACCGT

### S2. Crystal Assemblies and Arrangements

#### Locations of the Motifs

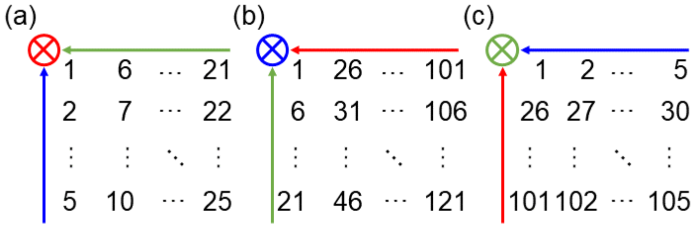


**Figure S3.** Locations of all the motifs in both 2T and 4T crystals in three different views. They are parallel to (a) green-blue plane, (b) red-green plane, and (c) blue-red plane. Similar to Figure 2(a), the arrow indicates the direction from 5' to 3' end. The circle crosshair means the arrow going into the page. In each view, there are five motif layers. In (a)-(c), the motifs are all at the bottom layer of such a view.

#### Connections of the Motifs

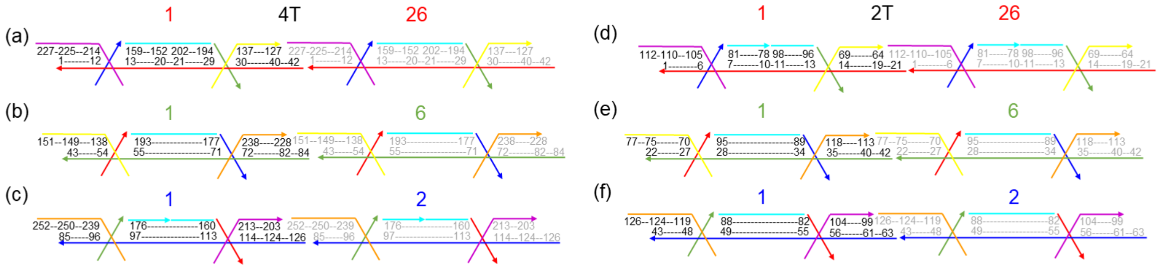


**Figure S4.** Connections of the motifs in red, green, and blue directions (via sticky-end association). After ligation, the neighboring 5' and 3' ends are merged. (a)-(c) on the left are for the 4T crystal, while (d)-(f) on the right are for the 2T crystal. The numbers above the strands are for the motifs, corresponding to those in Figure S3. The numbers on the strands are for the nt, which are the same as in Figures S1(b) or S2(b). To distinguish the motif on the left (motif number 1) and the one on the right (motif number 26, 6 or 2) in each panel, the nt labels are in different color. The left side is black, and the right side is gray.

Figure S3 illustrates where each motif is located, while the detailed connection is depicted in Figure S4. As an example for the connection, all the directions are shown as the connection between motif number 1 and its neighbors (number 26, 6 or 2, as also shown in Figure S3). The same connection would propagate in the same direction. For instance, the connection between 26 and 51 along the red direction is the same as between 1 and 26. Similarly, the link between 6 and 31 along the red is equivalent to between 1 and 26.

#### Outer Cross-Sectional Planes

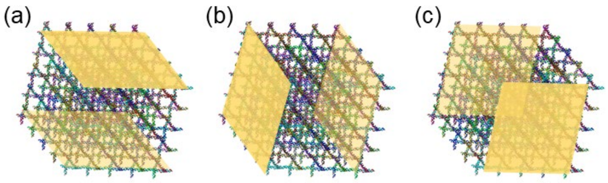


**Figure S5.** Indication of all 6 outer cross-sections, highlighted in gold color. The 4T crystal is the same as in Figure 1(d). (a) Top (dark gold) and bottom (light gold) cross-section planes. (b) Left (dark gold) and right (light gold) cross-section planes. (c) Front (dark gold) and back (light gold) cross-sections.

When applying the force plane, the holding plane is exerted on the nt which are located on the outer surface of the bottom cross-sectional plane. Similarly, the pulling plane is enforced on the nt which are on the outer surface of the top cross-sectional plane (Figure S5(a)). Since the red is the loading direction, Figure S3(a), Figure S4(a), and Figure S4(d) are used. The holding plane is exerted on motif number 1 to 25. In each motif, only one nt will be chosen. That is number 13 for the 4T crystal and 7 for the 2T crystal. Along this idea, the pulling plane is enforced on motif number 101 to 125, and on nt number 29 for the 4T crystal and 13 for the 2T crystal. Because we excluded the protruding part of the strand (*i.e.*, out of the bottom holding and the top pulling planes), there are only 185 bases per strand along the pulling direction rather than 210 in the 4T crystal. In the 2T crystal, it is 91 instead of 105.

### S3. Supporting Simulation Snapshots

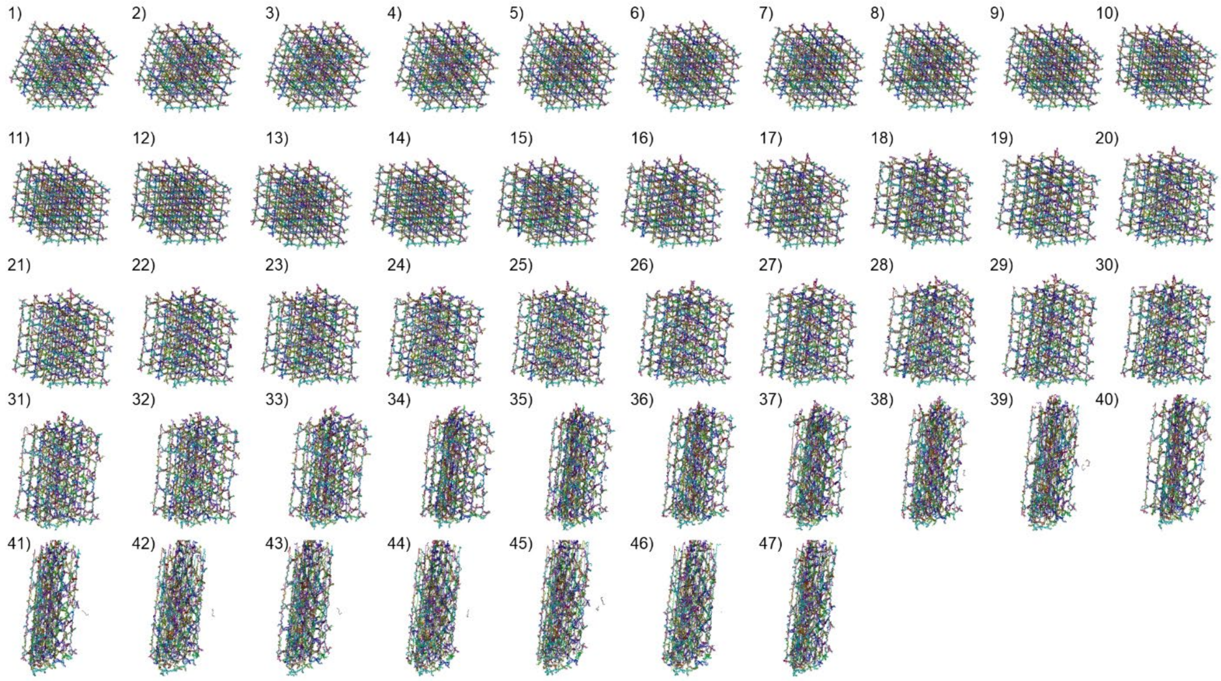

Figure S6. 47 snapshots in the deformation process of the 4T crystal in the MD simulations. These correspond to the data points in Figure 3(a), including the cross marks and the solid dots. Some of them (1, 3, 9, 24, and 40) are presented in Figure 2(b)-(f) and shown as the solid dots in Figure 3(a).

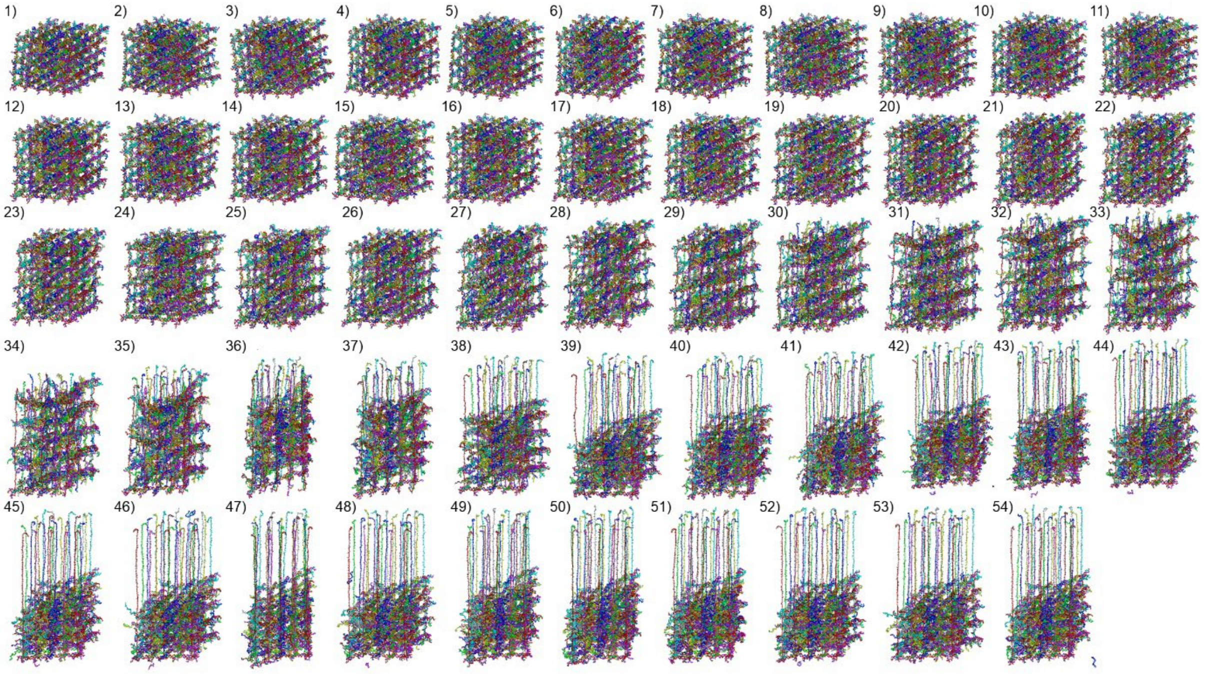

Figure S7. 54 snapshots in the deformation process of the 2T crystal in the MD simulations. These correspond to the data points in Figure 5(f) (both cross marks and solid dots). Some of them (1, 5, 25, and 44) are presented in Figure 5(b)-(e) and as the solid dots in Figure 5(f). The cross-sectional planes formed by the two non-loaded directions started to fail from the loaded direction in 30. In 39, the cross-sectional planes collapse completely.

### S4. Supporting Processed Simulation Data

#### Poisson's ratio of the 4T crystal

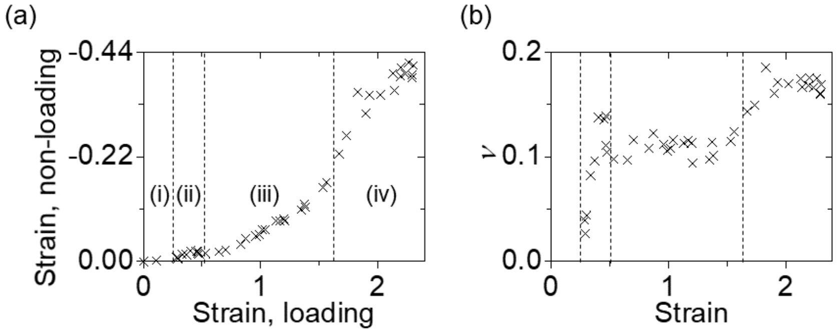


**Figure S8.** (a) Strain of averaged length in the non-loading (green and blue) directions vs. strain in the loading (red) direction. Negative value means contraction, whereas positive indicates elongation or expansion. (b) Poisson's ratio ($v$) with respect to the strain in the loading (red) direction. The stages are the same as in Figure 3.

As the 4T crystal extended in the loading direction, the non-loading directions contracted. In stage (i), the non-loading directions were subject to thermal fluctuation as the crystal was not on tension. Visible contraction started in stage (ii) and increased gradually in stage (iii). These indicate the initial development of the tension and the progressive loading on dsDNA. In the last stage, the crystal contracted further per unit elongation in the loading direction. This should be due to the pleated cross-sectional planes. They had little resistance against contraction after pleating.

Poisson's ratio ($v$) describes the relative changes in two orthogonal dimensions in a material under elastic stretch or compression.[1,2]

$$v = -\frac{\Delta y/y}{\Delta x/x} = -\frac{\varepsilon_y}{\varepsilon_x} \tag{S4.1}$$

$\varepsilon_x$ and $\varepsilon_y$ are strain in $x$ and $y$ directions, respectively. The value is between -1 and 0.5 for isotropic materials. With a positive Poisson's ratio, the two directions react in an opposite way. For example, $x$ is compressed to shrink, and then $y$ would expand. This is the most common scenario. A Poisson's ratio of $v = 0$ indicates the independence between the two directions. Whether $x$ direction is subjected to extension or compression, $y$ would not have any reaction. When the Poisson's ratio is negative, the two directions either expand or shrink together. This type of behavior is termed auxetics.[3] The 4T crystal has positive values of Poisson's ratio. Its non-loading directions always shrink upon extension in the red direction. In stage (ii), Poisson's ratio went from 0 to ~0.1 when contraction started. In stage (iii), Poisson's ratio remained at ~0.1. As the tensile forces were drastic in stage (iv), the ratio increased up to slightly below 0.2. The results are in good alignment with the observation of the simulation snapshots.

### S5. Nanoindentation Concepts and Verifications

#### Flat and topological designs

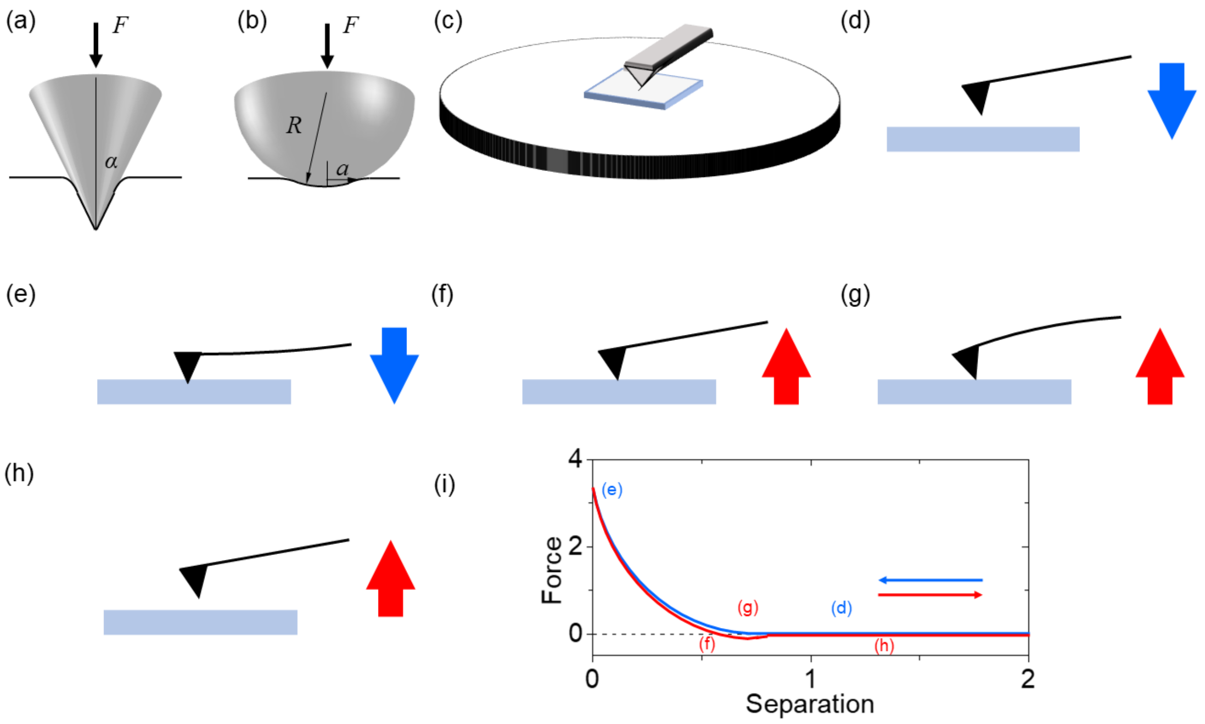


**Figure S9.** Two common nanoindentation models: (a) conical and (b) spherical. The key factor in the conical model is the half angle of the indenter, $\alpha$. Important parameters in the spherical model are radius $R$ of the indenter and the radius of indented cross-section, $a$. The indentation depth $\delta$ is not shown in the schematics. (c) AFM probe indenting a DNA crystal on a mica surface. The liquid environment was not drawn for simplicity. The detailed process is depicted in (d) through (h). (d)-(h) Blue and red mean approaching and retracting, respectively. This notation is the same as in Figure 4(c). (d) Approaching the crystal surface. (e) Reaching the maximum indenting force. (f) Retracting the probe. (g) Reaching the maximum adhesive force. (h) Back in the liquid. (i) General force trend for the process depicted in (d)-(h).

The expressions of the indentation force $F$ in the conical and spherical models are as follows.[4,5]

$$F = \frac{2}{\pi}\frac{E}{(1-\nu^2)}\delta^2 \tan\alpha \qquad \text{(S5.1)}$$

$$F = \frac{4}{3}\frac{E}{(1-\nu^2)}\delta^{3/2}\sqrt{R} \qquad \text{(S5.2)}$$

$E$ is the Young's modulus of the material being tested, $\nu$ is the Poisson's ratio, and $\delta$ is the indentation depth. The conical model (S5.1) is often used for deep indentation. The depth must be much greater than the tip radius if it is not super sharp. The spherical model (S5.2) is for shallow indentation with $\delta \leq R$. In general, the indentation depth should be at least 5 nm. We tested several PDMS samples to verify our system. The tip radius $R$ was around 10 nm. The indentation depth was similar to the tip radius, and thus, spherical model was used. The results were 2 – 6 MPa, which agrees well with the previous publications.[6-8]

### S6. References


1 Gibson, L. J. & Ashby, M. F. *Cellular Solids: Structure and Properties*. (Cambridge University Press, 1999).

2 Li, R., Chen, H. & Choi, J. H. Auxetic Two-Dimensional Nanostructures from DNA. *Angewandte Chemie* **60**, 7165-7173 (2021).

3 Chan, N. & Evans, K. E. Indentation Resilience of Conventional and Auxetic Foams. *Journal of Cellular Plastics* **34**, 231-260 (1998).

4 Harding, J. & Sneddon, I. in *Mathematical Proceedings of the Cambridge Philosophical Society.* 16-26 (Cambridge University Press).

5 Derjaguin, B. V., Muller, V. M. & Toporov, Y. P. Effect of Contact Deformations on the Adhesion of Particles. *Journal of Colloid and Interface Science* **53**, 314-326 (1975).

6 Wang, Z., Volinsky, A. A. & Gallant, N. D. Nanoindentation Study of Polydimethylsiloxane Elastic Modulus Using B Erkovich and Flat Punch Tips. *Journal of Applied Polymer Science* **132** (2015).

7 Charitidis, C. A. Nanoscale Deformation and Nanomechanical Properties of Polydimethylsiloxane (PDMS). *Industrial & Engineering Chemistry Research* **50**, 565-570 (2011).

8 Deuschle, J. K., Buerki, G., Deuschle, H. M., Enders, S., Michler, J. & Arzt, E. In situ Indentation Testing of Elastomers. *Acta Materialia* **56**, 4390-4401 (2008).
